# Newcastle Disease Virus Vector-Based SARS-CoV-2 Vaccine Candidate AVX/COVID-12 Activates T Cells and Is Recognized by Antibodies from COVID-19 Patients and Vaccinated

**DOI:** 10.1101/2024.03.01.582987

**Authors:** Alejandro Torres-Flores, Luis Alberto Ontiveros-Padilla, Ruth Lizzeth Madera-Sandoval, Araceli Tepale-Segura, Julián Gajón-Martínez, Tania Rivera-Hernández, Eduardo Antonio Ferat-Osorio, Arturo Cérbulo-Vázquez, Lourdes Andrea Arriaga-Pizano, Laura Bonifaz, Georgina Paz-De la Rosa, Oscar Rojas-Martínez, Alejandro Suárez-Martínez, Gustavo Peralta-Sánchez, David Sarfati-Mizrahi, Weina Sun, Héctor Elías Chagoya-Cortés, Peter Palese, Florian Krammer, Adolfo García-Sastre, Bernardo Lozano-Dubernard, Constantino López-Macías

## Abstract

Several effective vaccines for severe acute respiratory syndrome coronavirus 2 (SARS-CoV-2) have been developed and implemented in the population. However, the current production capacity falls short of meeting global demand. Therefore, it is crucial to further develop novel vaccine platforms that can bridge the distribution gap. AVX/COVID-12 is a vector-based vaccine that utilizes the Newcastle Disease virus (NDV) to present the SARS-CoV-2 spike protein to the immune system. This study analyses the antigenicity of the vaccine candidate by examining antibody binding and T-cell activation in individuals infected with SARS-CoV-2 or variants of concern (VOCs), as well as in healthy volunteers who received coronavirus disease 2019 (COVID-19) vaccinations. Our findings indicate that the vaccine effectively binds antibodies and activates T-cells in individuals who received 2 or 3 doses of BNT162b2 or AZ/ChAdOx-1-S vaccines. Furthermore, the stimulation of T-cells from patients and vaccine recipients with AVX/COVID-12 resulted in their proliferation and secretion of interferon-gamma (IFN-γ) in both CD4+ and CD8+ T-cells. In conclusion, the AVX/COVID-12 vectored vaccine candidate demonstrates the ability to stimulate robust cellular responses and is recognized by antibodies primed by the spike protein present in SARS-CoV-2 viruses that infected patients, as well as in the mRNA BNT162b2 and AZ/ChAdOx-1-S vaccines. These results support the inclusion of the AVX/COVID-12 vaccine as a booster in vaccination programs aimed at addressing COVID-19 caused by SARS-CoV-2 and its VOCs.

## Introduction

Severe acute respiratory syndrome coronavirus 2 (SARS-CoV-2) and its variants of concern have caused billions of infections and millions of deaths worldwide since 2020. Despite the incredible speed of coronavirus disease 2019 (COVID-19) vaccine development since 2020, there have been more than 7 million deaths from COVID-19, with an estimated excess mortality of 28.4 million deaths (1,2). The unequal distribution of vaccines globally, especially impacting low- and middle-income countries (LMICs), underscores the urgency for further advancements in new vaccine platforms to address the disparity in vaccine access (3).

The AVX/COVID-12 vaccine candidate is a Newcastle disease virus (NDV) LaSota vector-based vaccine that expresses the stabilized form of the SARS-CoV-2 spike protein. The stabilization is achieved by introducing six prolines (HexaPro-S), ensuring the protein remains in its closed conformation (4). This vaccine has demonstrated safety, immunogenicity, and protective efficacy in preclinical models (5,6), and has the advantage of being produced in embryonated egg industrial facilities similarly to influenza vaccine production which is cheap and widely available (7). In a phase I clinical trial, the AVX/COVID-12 vaccine demonstrated safety, good tolerability, and immunogenicity in volunteers (8), supporting further clinical development of the vaccine. However, high rates of infections in the population and the availability of approved vaccines in Mexico made the development of a phase II placebo-controlled clinical trial unethical. To provide evidence supporting the testing of this vaccine as a booster dose, we analysed the capacity of the AVX/COVID-12 vaccine to bind antibodies and stimulate T-cell responses from individuals previously infected or vaccinated with approved vaccines.

## Material and methods

### Study groups

Samples were collected from 41 volunteers who read and signed the informed consent for this study. The subjects were categorized into five groups: patients with COVID-19 in the acute phase of the disease (7 days after hospital admission; n=12) and a subgroup infected with the Omicron variant (Omicron peak, December 2021; n=10) who reported no previous infection or vaccination, patients who had recovered from COVID-19 during the first peak of the pandemic (June-December 2020; n=6), healthy volunteers vaccinated with two doses of Pfizer-BioNTech BNT162b2 (4 to 8 months after the second dose; n=6) and volunteers with two doses of Pfizer-BioNTech BNT162b2 boosted with AstraZeneca AZ/ChAdOx-1-S (4 to 8 months after the second dose; n=7).

### Serum collection

Venous blood samples were obtained from participants, and these samples were collected using two ethylenediaminetetraacetic acid (EDTA) tubes and one red tube (BD Vacutainer tubes, Franklin Lakes, NJ, USA) following standard phlebotomy procedures. After collection, all blood samples and their derivatives were processed in a biosafety level (BSL)-2 laboratory, with the use of appropriate personal protective equipment and safety precautions. Serum isolation involved centrifuging venous blood (collected in red tubes) at 2,000 x *g* for 10 minutes to separate the serum. The resulting serum was carefully extracted from the upper portion of the tube, aliquoted, and subsequently stored at −20 °C until needed.

### Peripheral blood mononuclear cell (PBMC) isolation

PBMCs were isolated from venous blood collected in EDTA tubes (BD vacutainer tubes, Franklin Lakes, NJ, USA). Within 4 hours (h) of collection, PBMC isolation was conducted by density-gradient sedimentation of whole blood diluted at a 1:2 ratio in phosphate-buffered saline (PBS) at room-temperature. The diluted blood was layered over an appropriate volume of room-temperature Lymphoprep (Serumwerk Bernburg AG, DEU; cat. 07851). Then, the PBMCs were recovered, cryopreserved in a medium consisting of 10% dimethyl sulfoxide (DMSO; Sigma Aldrich, St. Louis, MO, USA) and 90% heat-inactivated fetal bovine serum (FBS; GIBCO, California, USA; cat. 11560636), and stored at −80 °C until use.

### Vaccine manufacturing

The AVX/COVID-12 vaccine was cultivated in 10-day-old specific pathogen-free (SPF) chicken embryos through inoculation into the allantoic cavity, using 10^3.3^ 50% egg infectious dose (EID_50_)/0.1□mL of the production seed. The embryos were incubated for 72□h at 37 °C with 60-70% humidity. After incubation, the embryos were refrigerated for a minimum of 12 h, and the allantoic fluid (AF) was then aseptically harvested. The AF underwent clarification through filters, was concentrated by a factor of 10X using 300 kDa cassettes, and was subsequently diluted in 20 volumes of PBS. The resulting AF was frozen and stored at −70°C until needed. The experimental AVX/COVID-12 vaccine, along with NDV-V (empty vector), was provided frozen in 2 mL vials. The vaccine was produced in Avimex’s good manufacturing practice facilities in Mexico City.

### Antibody analysis by enzyme-linked immunosorbent assay (ELISA)

The analysis of specific anti-spike IgG antibodies was conducted using ELISA. Each plate was coated with NDV-V, AVX/COVID-12, or the ancestral receptor binding domain (RBD) of SARS-CoV-2 at a concentration of 10 μg/mL. Serum samples, prediluted at 1:40, were transferred to the plate (200 μL), and then 1:2 serial dilutions were performed. The plates were incubated at 37 °C for 60 minutes and subsequently washed with tris-buffered saline (TBS)-Tween 20 buffer. A 1:4000 dilution of anti-human IgG coupled to horseradish peroxidase (HRP; MyBioSource’s, San Diego, USA; cat. MBS440121) was added to the plates, followed by an incubation at 37 °C for 60 minutes. To stop the reaction, 2.5 N sulfuric acid was added, and the optical density (OD) was read at 450 nm using the Epoch Microplate Spectrophotometer (BioTek Laboratories, Seattle, Washington, USA) within 10 minutes after adding the stop solution. The antibody titre was determined as three times the value of the negative controls, and titres were represented as −Log2×40.

### T-cell proliferation assay

PBMCs were washed twice with 1X PBS to eliminate the cryopreservation medium, and the cell button was resuspended and adjusted to 5 x 10^6^ cells/mL. Subsequently, proliferation staining was carried out using the CellTrace Violet reagent (Invitrogen, Massachusetts, USA; cat. C34557) at 37 °C for 20 min. Every 5 min, the cells were vortexed to ensure uniform staining. Following staining, the cells were washed with 1X PBS and then with Roswell Park Memorial Institute (RPMI) 1640 culture medium (GIBCO, Thermo Fisher, Massachusetts, USA) without supplementation. After this incubation period, the stained cells were harvested, washed with fresh medium, counted, and cultivated in 96-well plates (5 x 10^5^ cells/well). Subsequently, they were stimulated separately with 10 μg of NDV-V, AVX/COVID-12 or 30 nM of a mixture of 10-15 amino acid long peptides derived from the SARS-CoV-2 spike protein (PepTivator, Miltenyi Biotec, North Rhine-Westphalia, Germany). In parallel, stained and unstimulated cells were cultivated at the same concentration as the controls. After this step, the cells were cultured in RPMI 1640 with 10% FBS and incubated for 72 h at 37 °C. The cells were harvested and surface-stained with anti-CD3 mAb APC (BioLegend, San Diego, California, USA; cat. 344812), anti-CD4 mAb APC-Cy7 (BioLegend, San Diego, California, USA; cat. 317418), and anti-CD8 mAb FITC (BioLegend, San Diego, California, USA; cat. 344704) for PBMCs from critically ill patients hospitalized during the first wave of COVID-19. For PBMCs from the other study groups, cells were stained with anti-CD3 mAb AF647 (BioLegend, San Diego, California, USA; cat. 300322), anti-CD4 mAb APC/Cy7 (BioLegend, San Diego, California, USA; cat. 317418), and anti-CD8 mAb PerCP/Cy5.5 (BioLegend, San Diego, California, USA; cat. 300924). The staining mixture was incubated for 15 minutes at room-temperature in the dark, followed by an additional washing step with cell staining buffer. Subsequently, the cells were fixed and permeabilized using Cytofix/Cytoperm for 20 minutes at room-temperature. After centrifugation, Perm/Wash was added. Intracellular staining was then performed using anti-interferon (IFN)-γ mAb PE (BioLegend, San Diego, California, USA; cat. 506507) for critically ill patients hospitalized and anti-IFN-γ mAb FITC (BioLegend, San Diego, California, USA; cat. 502506) for PBMCs from the other study groups in Perm/Wash solution (BD, San Jose, USA) for 30 minutes at room-temperature in the dark. After washing with BD Perm/Wash buffer (BD, San Jose, USA), the cells were resuspended in PBS and kept at 4 °C in the dark until acquisition and analysis. Flow cytometric analysis was performed using the FACS Canto II flow cytometer (BD Biosciences, New Jersey, USA), and the data were analysed using FlowJo V10 software (Fig. 1S).

**Figure 1.**
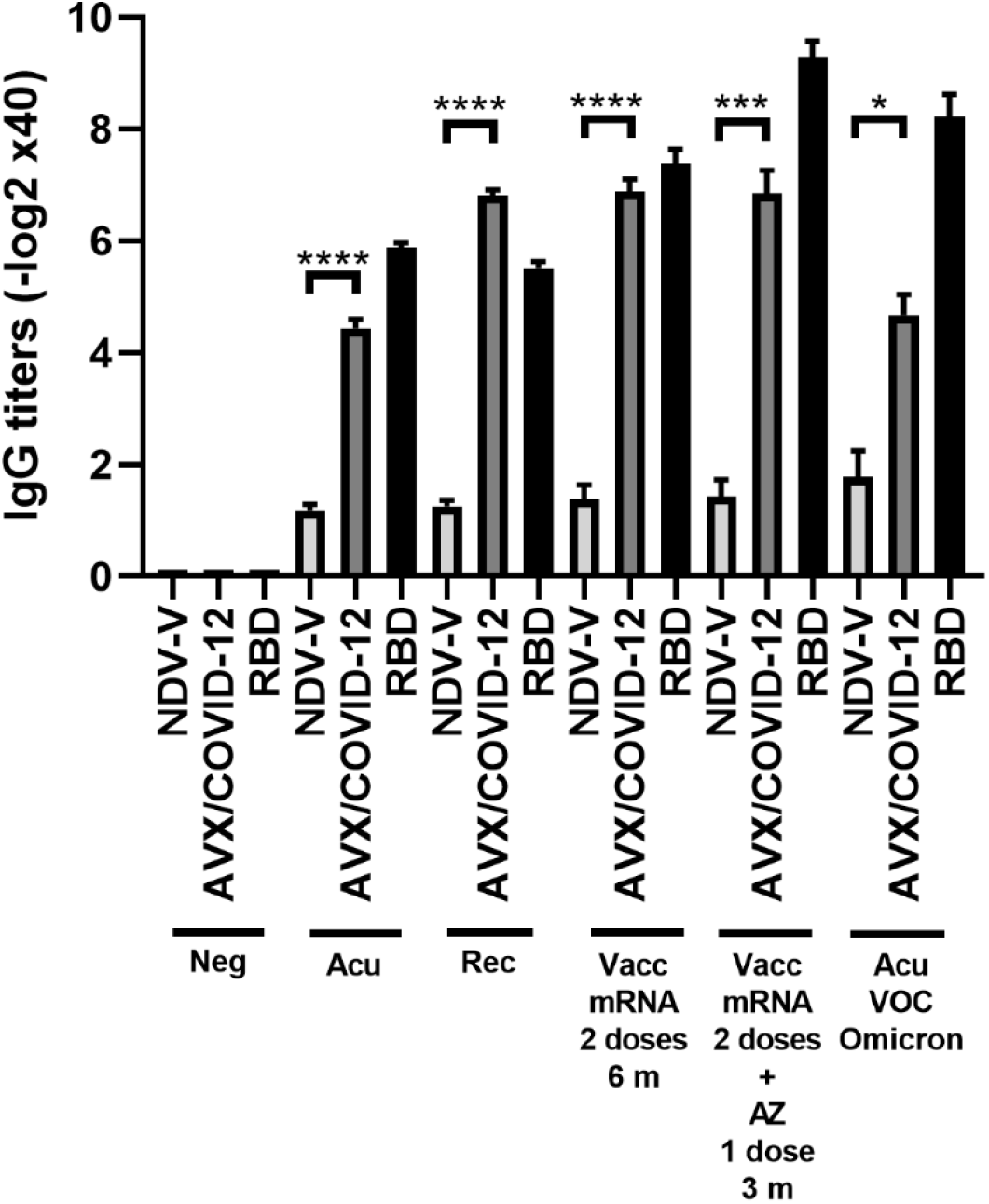
Binding of serum IgG from COVID-19 patients and vaccinated individuals to the AVX/COVID-12 vaccine and the Receptor-Binding Domain (RBD) of the spike protein of SARS-CoV-2. ELISAs were conducted to assess the binding of the AVX/COVID-12 vaccine and the receptor binding domain (RBD) to serum IgG antibodies from patients in the acute phase of COVID-19 (Acu) (n=12) and a subgroup with the Omicron variant (Acu VOC Omicron) (n=10), as well as individuals who had recovered from the disease (Rec) (n=6). Additionally, samples were collected from individuals vaccinated with two doses of the BNT162b2 Pfizer vaccine after 6 months of the second dose (Vacc mRNA) (n=6) and those who received two doses of Pfizer and were boosted with the AZ (AZ/ChAdOx-1-S) vaccine after 3 months of the third dose (n=7). Control groups included NDV-V (NDV LaSota virus empty vector) and AVX/COVID-12 (NDV expressing SARS-CoV-2 S protein and RBD). Kruskal Wallis. p<0.05*, p<0.01**, p<0.001*** and p<0.0001****.

### Statistical analysis

Descriptive statistics were employed to analyse the variables of interest. For discrete or continuous quantitative variables, the mean, median, and standard deviation were determined as appropriate for each case. The choice between parametric or non-parametric statistics was made based on the distribution and variance of the data. Regarding the analysis of antibody titres and cell percentages, calculations were performed for frequencies, means, or distributions. Student’s t-test or Mann-Whitney U test was used for two categories, while 2-way ANOVA and Bonferroni post-test or Kruskal-Wallis were employed for the comparison of more than two groups. The results were processed using Excel, the GraphPad Prism analysis program, and STATA v.14 software.

## Results

### Analysis of antibody response for antigenicity evaluation of the AVX/COVID-12 vaccine

To evaluate the antigenicity of AVX/COVID-12, we performed *in vitro* tests with antibodies induced after infection or vaccination against COVID-19. Utilizing ELISA, we assessed the binding of these antibodies to the AVX/COVID-12 vaccine, NDV-V, and recombinant RBD (Fig. 1).

We observed specific AVX/COVID-12 IgG antibody titres ranging from 4 to 7 logarithmic units base 2 (log2 units) in sera from both acute and recovered SARS-CoV-2 infected patients. Notably, sera from hospitalized patients during the Omicron wave, who had not received prior COVID-19 vaccination in Mexico, also exhibited binding to AVX/COVID-12, similar to sera from patients infected with other VOCs. In vaccinated individuals, we observed statistically significant differences with higher titres, ranging from 7 to 9. In contrast, NDV-V, which lacks SARS-CoV-2 spike protein expression, exhibited only marginal binding with a titre of 1. Using RBD as a positive control for coating the plates, we found no statistical differences between antibody titres binding to RBD or the AVX/COVID-12 vaccine. These results suggest that the spike protein expressed in the AVX/COVID-12 vaccine enables the binding of IgG antibodies triggered by the spike protein expressed by the SARS-CoV-2 variant that infected patients, as well as by the BNT162b2 and AZ/ChAdOx-1-S vaccines.

### AVX/COVID-12 vaccine elicits T-cell proliferation and IFN-γ production in patients and COVID-19 vaccinated volunteers

Next, we investigated whether the AVX/COVID-12 vaccine activates T-cells in individuals who had COVID-19 or were vaccinated with anti-COVID-19 vaccines. We stimulated PBMCs from patients (Fig. 2) and vaccinated volunteers (Fig. 3) with AVX/COVID-12 or the NDV-V empty vector. As a positive control, we utilized a spike protein peptide cocktail mixture of 10-15 amino acid peptides (PepTivator S) (Fig. 1S).

**Figure 2.**
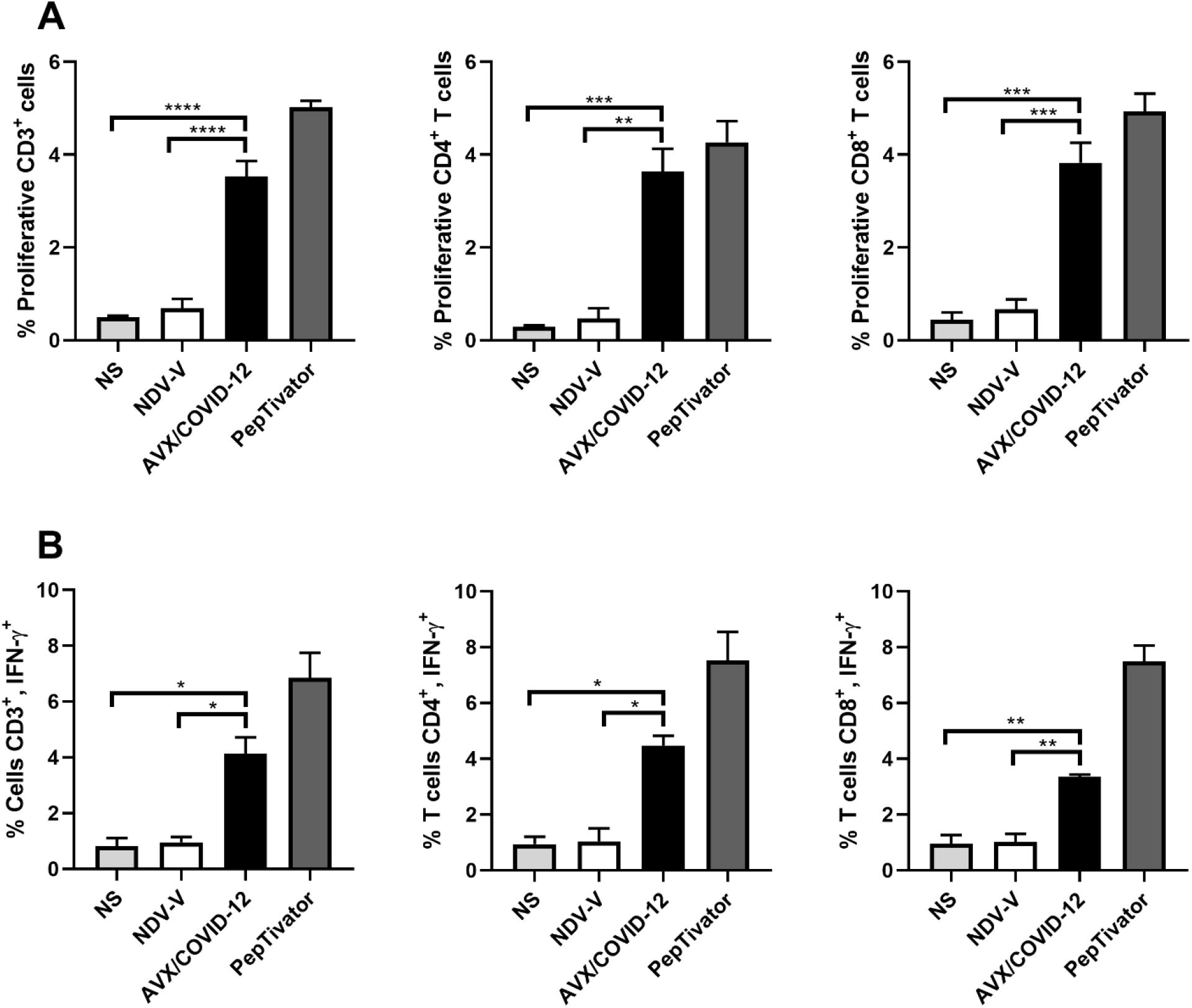
AVX/COVID-12 Vaccine Induces Proliferation and IFN-γ Response in T Cells from Acute COVID-19 Patients. Proliferation response (A) or the percentage of IFN-γ+ (B) in T cells from acute COVID-19 patients (ancestral first wave) was assessed following stimulation with NDV-V (empty vector of NDV LaSota virus), AVX/COVID-12, or PepTivator (S1 and S2 peptides) for 72 hours. Flow cytometry analysis encompassed individual evaluations of CD4+ and CD8+ responses in single, FSClow SSClow CD3+ cells. NS (Not Stimulated). n=3, ANOVA, Tukey. p<0.05*, p<0.01**, p<0.001*** and p<0.0001****.

**Figure 3.**
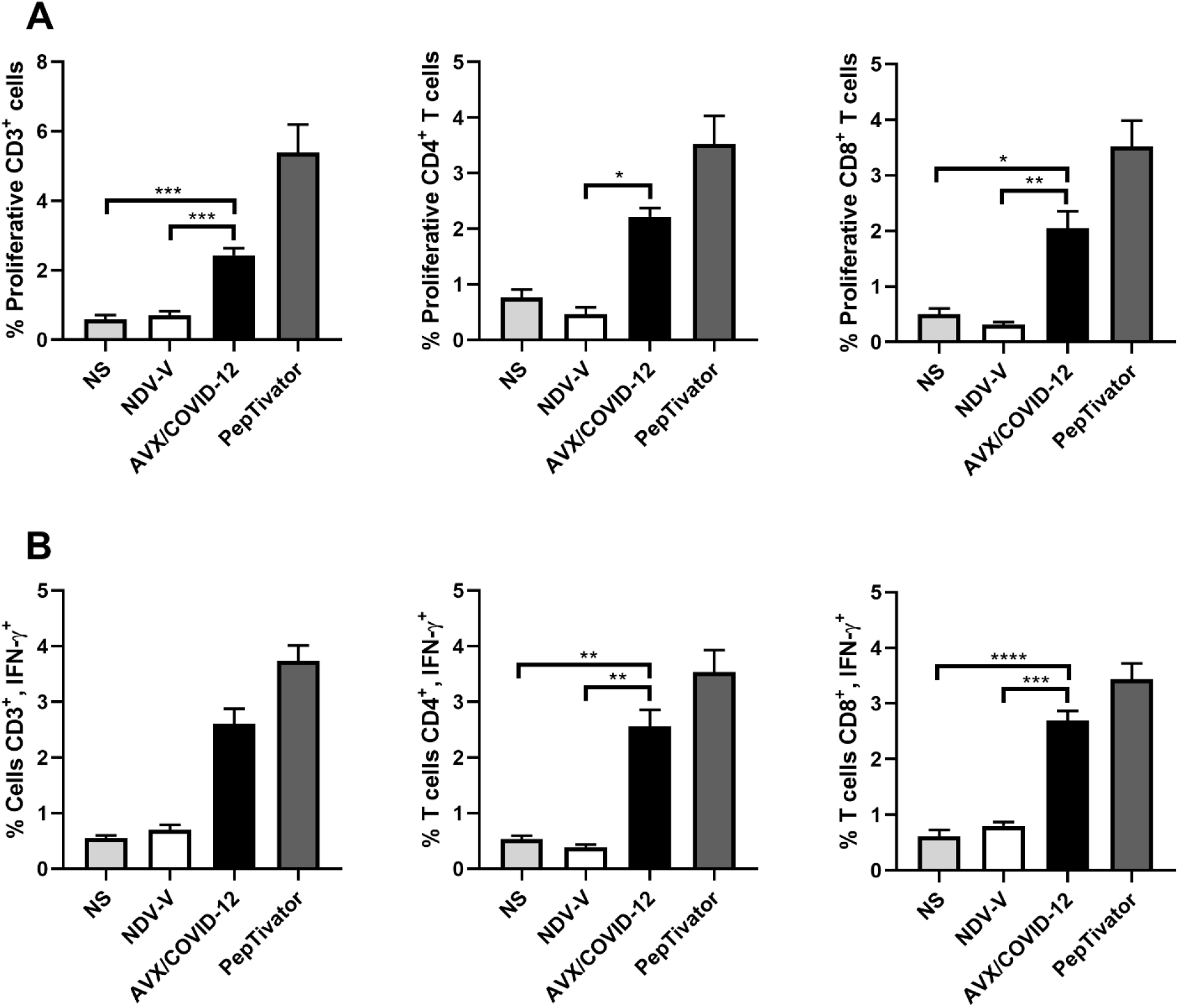
AVX/COVID-12 Vaccine Elicits Proliferation and IFN-γ Response in T Cells from Individuals Vaccinated with mRNA BNT162b2. Proliferation response (A) or percentage of IFN-γ+ (B) in T cells stimulated with NDV-V (empty vector of NDV LaSota virus), AVX/COVID-19 vaccine, or PepTivator (S1 and S2 peptides) for 72 hours. Flow cytometry analysis includes single, FSClow SSClow CD3+ cells, and individual responses of CD4+ and CD8+, respectively. NS (Not Stimulated). n=6, Kruskal Wallis or ANOVA, Tukey. p<0.05*, p<0.01**, p<0.001*** and p<0.0001****.

We observed that the AVX/COVID-12 vaccine induced specific proliferation and intracellular IFN-γ production in both critical COVID-19 patients and vaccinated volunteers. In contrast, NDV-V induced a basal response similar to that of non-stimulated cells (Fig. 2A and 3A). The positive control, PepTivator, also triggered strong specific cell proliferation and IFN-γ intracellular production, surpassing AVX/COVID-12 vaccine by 1-2%.

The production of intracellular IFN-γ in total CD3^+^ T cells from patients was higher than in vaccinated volunteers (p = 0.0169 and p = 0.0009). The same trend was observed for CD4^+^ T-cells (p = 0.0011 and p = 0.0437) and CD8^+^ T-cells (p = 0.0005 and p = 0.0035). Additionally, we observed an increase in the proliferation response in T-cells from critically ill acute patients compared to those vaccinated with two doses of the BNT162b2 vaccine (Fig. 2A and 3A).

With a vaccine booster, one would anticipate a substantial increase in both antibody and cellular responses, potentially ranging from 10 to 25 times greater (9,10). Therefore, we examined the cellular response in PBMCs from volunteers who had received two doses of BNT162b2 and were subsequently boosted with the AZ/ChAdOx-1-S vaccine, stimulated with the AVX/COVID-12 vaccine (Fig. 4). We detected specific proliferation and intracellular production of IFN-γ induced by the AVX/COVID-12 vaccine in CD3+, CD4+, and CD8+ T-cells, while the NDV-V empty vector induced basal activation similar to that of non-stimulated cells. Nevertheless, the response in volunteers who received a booster dose with the AZ/ChAdOx-1-S vaccine is comparable to that of volunteers who received only two doses of the BNT162b2 vaccine, with a slight tendency to be lower when compared to the response observed in T-cells from critically ill COVID-19 patients.

**Figure 4.**
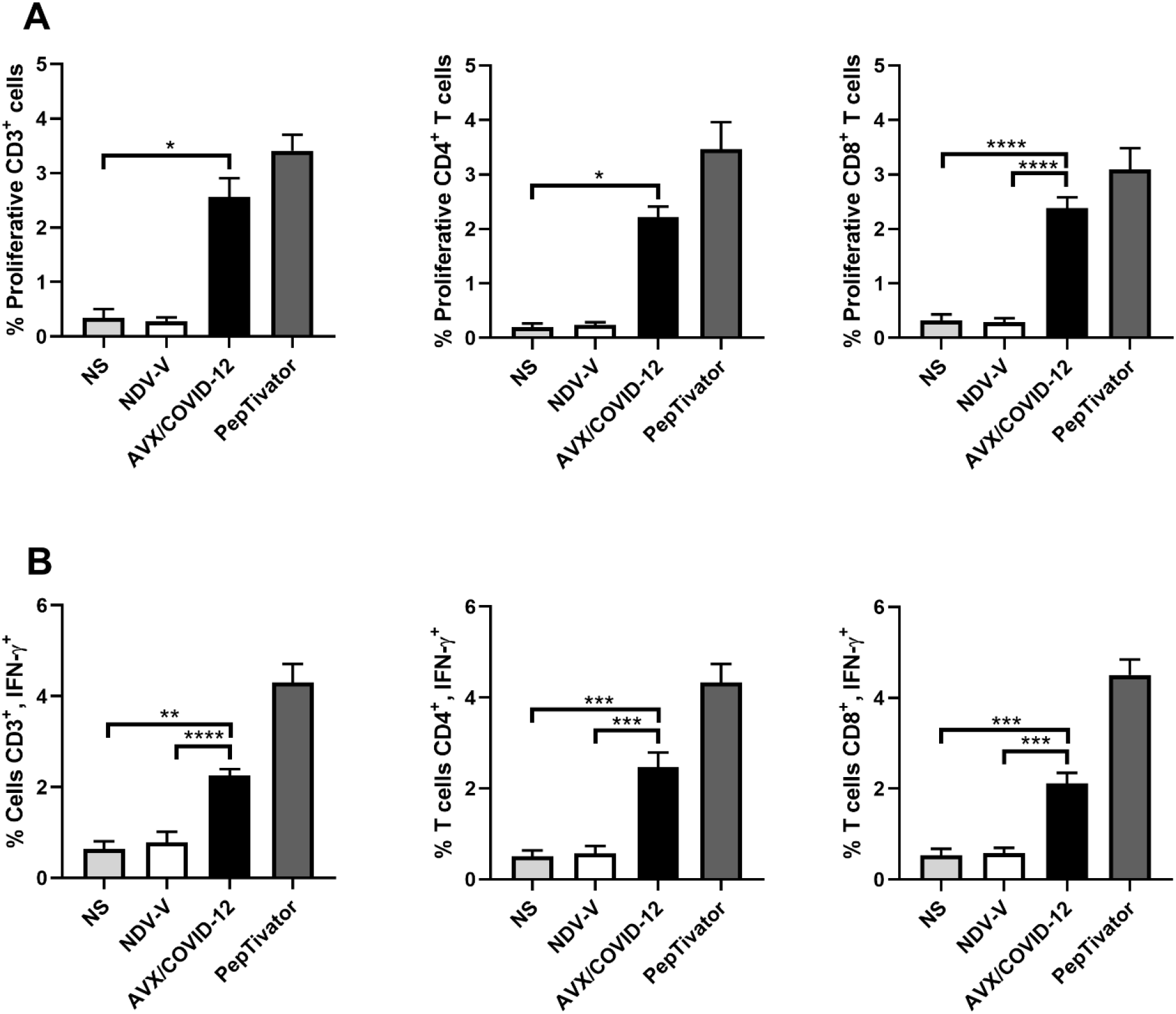
AVX/COVID-12 Vaccine Induces Proliferation and IFN-γ Response in T Cells from Individuals Vaccinated with mRNA BNT162b2 and Boosted with AZ/ChAdOx-1-S Vaccine. Proliferation response (A) or percentage of IFN-γ+ (B) in T cells stimulated with NDV-V (empty vector of NDV LaSota virus), AVX/COVID-12, or PepTivator (S1 and S2 peptides) for 72 hours. Flow cytometry analysis includes single, FSClow SSClow CD3+ cells and individual response of CD4+ and CD8+ respectively. NS (no stimulated). n=6, Kruskal Wallis. p<0.05*, p<0.01**, p<0.001*** and p<0.0001****.

## Discussion

During the SARS-CoV-2 pandemic, there was a pressing need for accelerated technological advancements in the development of new vaccine platforms. mRNA vaccines and viral vectors emerged as frontrunners, receiving emergency use approvals, and demonstrating their efficacy in alleviating hospital saturation and reducing deaths due to COVID-19. However, to achieve broader vaccination coverage, especially in LMICs, there is a demand for vaccine candidates that are not only effective but also affordable and easily transportable. The NDV viral vector platform has proven to be a safe and innocuous vector (5,6,11). It is easily developed for use, showing high effectiveness in oncolytic treatment and preclinical models as a live vaccine vector (12,13). This makes it a promising candidate for meeting the vaccine needs in diverse settings.

The use of the NDV-HXP-S (AVX/COVID-12-HEXAPRO) vector technology has facilitated the development of the AVX/COVID-12 vaccine candidate, demonstrating effectiveness in eliciting neutralizing antibodies and an IFN-γ^+^ T-cell response crucial for combating SARS-CoV-2 infection (5,7).

Due to the updated COVID-19 vaccine distribution inequities affecting mainly LMICs, it is crucial to gather evidence on the efficacy of currently available vaccines and vaccine candidates. This should focus on assessing the capacity of the spike protein from the original Wuhan-1 strain to generate immunity against emerging VOCs such as Omicron and its subvariants.

If the epitopes constituting the spike protein of the SARS-CoV-2 or those from the first-generation vaccines (BNT162b2 and AZ/ChAdOx-1-S) are efficiently expressed in the NDV vector, we anticipate the development of antibodies targeting similar epitopes. These antibodies would play a crucial role in protecting the host from the development of severe and critical pathology (14–16).

One approach to evaluate the capacity of a vaccine candidate in eliciting immune responses in humans involves *ex vivo* tests assessing its antigenicity. This process entails examining the candidate’s interaction with pre-existing antibodies developed in patients who have recovered from the targeted pathology or through the recognition and activation of the cellular immune response.

We observed the binding of antibodies to the spike protein in comparison to the empty vector NDV-V or the RBD in different study groups during and after infection or post-vaccination with second and third doses. Anti-spike IgG titres showed an increase of 4 to 6 log2 units across all study groups (Fig. 1), indicating similar humoral responses between the two vaccination schemes. In this context, the design of the vaccine protein S in BNT162b2 and AVX/COVID-12 shares the prefusion-stabilized structure (6). However, there are differences in the substitutions made to maintain the structure of the molecule, involving two prolines in BNT162b2 and six prolines in the S2 subunit of AVX/COVID-12 (6,17). This suggests antibodies are targeting conformational epitopes allowing both candidates to potentially share structural epitopes. When comparing the antigenicity against antibodies developed with a third dose of the AZ/ChAdOx-1-S vaccine, where the structure of the protein S resembles the native one (18), we observed similar activation (without significant differences, p > 0.9) compared to the complete scheme with BNT162b2 alone. Therefore, all three designs appear to induce similar repertoires of antibodies against SARS-CoV-2.

Moreover, upon analysing the antibody response in patients during the acute phase and those who have recovered from COVID-19, we observed that recovered patients had 3 log2 units higher anti-spike IgG antibody titres than acute patients (p = 0.1533) (Fig. 1). In contrast, it has previously been reported that there are no significant differences in antibody titres among acute and recovered patients (19). However, differences in the quality of the response are observed, particularly in terms of opsonization, complement activity, and neutralizing capability (20). This is attributed to the significant reduction in the percentage and total numbers of T and B lymphocyte populations during severe acute COVID-19. This reduction reflects a noteworthy decrease in the development of germinal centres in lymph nodes, potentially impairing the affinity and repertoire of antibodies to various epitopes of the total spike protein (21,22). The increase of antibody titres in recovered patients suggests a gradual increase in the number of epitopes recognized by the immune system during the recovery phase from COVID-19.

In the design of vaccines or platforms, the evaluation of T-cell activation response has been a key aspect. For instance, in cases like the respiratory syncytial virus in infants or the pandemic influenza H1N1 virus, assays involving lymphoproliferation and IFN-γ secretion have been utilized as indicators of T-cell effector activity, providing valuable insights into the vaccine’s effectiveness (23,24). We analysed three pivotal scenarios during the pandemic: the cellular response in COVID-19 patients, emphasizing the effector response; the vaccination schemes, specifically 6 months after the second dose to ensure memory T-cell stimulation in peripheral blood; and 3 months after the third dose, exemplifying restimulation with a heterologous immunization schedule. The analysis revealed that, 72 h after stimulation, proliferation was induced with the vaccine in the case of patients (Fig. 3 and Fig. 4). This was expected, as despite a low percentage of T-cells in patients, they exhibited a robust effector response primarily targeting spike and nucleocapsid protein epitopes. Consequently, these T-cells were activated by AVX/COVID-12 and produced IFN-γ. Regarding vaccinated volunteers, notable differences were observed in both cases. Surprisingly, when comparing proliferation and interferon responses, they appeared to be similar between both doses (data not reported). This contradicts expectations, since with a vaccine booster, one would anticipate a substantial increase in both antibody and cellular responses, ranging from 10 to 25 times greater (9,10). However, since we are examining antigenicity, we can only illustrate the epitopes present in the vaccine. Therefore, upon immunization with the candidate, we anticipate the response to escalate. These observations are consistent with the identification of a substantial number of conserved epitopes within the spike protein, contributing to both humoral and cellular responses across ancestral SARS-CoV-2 and its variants of concern, as well as SARS-CoV, MERS-CoV, and seasonal coronaviruses (25,26). This may even extend to other coronaviruses within the *Orthocoronavirinae* genus (27).

Furthermore, considering the inoculation route (potential nasal administration in the case of AVX/COVID-12), it is expected to stimulate the development of both mucosal and systemic responses. These aspects will be further elucidated during the upcoming phases of the clinical trials for the AVX/COVID-12 vaccine candidate.

In conclusion, the AVX/COVID-12 vectored vaccine demonstrates the ability to stimulate robust cellular responses and is recognized by antibodies primed by the spike protein present in SARS-CoV-2 viruses that infected patients, as well as in the mRNA BNT162b2 and AZ/ChAdOx-1-S vaccines. These findings endorse the integration of the AVX/COVID-12 vaccine as a booster in vaccination initiatives designed to combat COVID-19 caused by SARS-CoV-2 and its VOCs.

## Conflict of Interest

The vaccine candidate used in this study was developed by faculty members at the Icahn School of Medicine at Mount Sinai including P.P., F.K. and A.G.-S. Mount Sinai is seeking to commercialize this vaccine; therefore, the institution and its faculty inventors could benefit financially. The Icahn School of Medicine at Mount Sinai has filed patent applications relating to SARS-849 CoV-2 serological assays (USA Provisional Application Numbers: 62/994,252, 63/018,457, 63/020,503 and 63/024,436) and NDV-based SARS-CoV-2 vaccines (USA Provisional Application Number: 63/251,020) which list F.K. as co-inventor. A.G.-S. and P.P. are a co-inventor in the NDV-based SARS-CoV-2 vaccine patent application. Patent applications were submitted by the Icahn School of Medicine at Mount Sinai. Mount Sinai has spun out a company, Kantaro, to market serological tests for SARS-CoV-2 and another company, CastleVax, to commercialize SARS-CoV-2 vaccines. F.K., P.P. and A.G.-S. serve on the scientific advisory board of CastleVax and are listed as co-founders of the company. F.K. has consulted for Merck, Seqirus, Curevac, and Pfizer, and is currently consulting for Gritstone, Third Rock Ventures, GSK, and Avimex. The F.K. laboratory has been collaborating with Pfizer on animal models of SARS-CoV-2. The A.G.-S. laboratory has received research support from GSK, Pfizer, Senhwa Biosciences, Kenall Manufacturing, Blade Therapeutics, Avimex, Johnson & Johnson, Dynavax, 7Hills Pharma, Pharmamar, ImmunityBio, Accurius, Nanocomposix, Hexamer, N-fold LLC, Model Medicines, Atea Pharma, Applied Biological Laboratories and Merck. A.G.-S. has consulting agreements for the following companies involving cash and/or stock: Amovir, Vivaldi Biosciences, Contrafect, 7Hills Pharma, Avimex, Pagoda, Accurius, Esperovax, Farmak, Applied Biological Laboratories, Pharmamar, CureLab Oncology, CureLab Veterinary, Synairgen, Paratus, Pfizer and Prosetta. A.G.-S. has been an invited speaker in meeting events organized by Seqirus, Janssen, Abbott, and AstraZeneca. P.P. has a consulting agreement with Avimex.

The live vaccine used in the study was developed by members of Avimex. Avimex filed patent applications with Mount Sinai and CONAHCYT. M.T., D.S.-M., C.L.-M., H.E.C.-C., F.C.-P., G.P.D.L., and B.L.-D. are named as inventors on at least one of those patent applications. The rest of the participants are employees of their corresponding institutions and declare no competing interests.

## Author Contributions

A.T.-F. Data curation, Formal Analysis, Investigation, Visualization, Writing – original draft, Writing – review & editing; L.A.O.-P. Formal Analysis, Investigation, Visualization, Writing – review & editing; R.L.M.-S. Data curation, Formal Analysis, Investigation, Methodology, Writing – review & editing; A.T.-S. Data curation, Writing – review & editing; J.G.-M. Data curation, Writing review & editing; T.R.-H. Data curation, Formal Analysis, Investigation, Supervision, Validation, Visualization, Writing – review & editing; E.A.F.-O. Data curation, Formal Analysis, Validation, Writing – review & editing; A.C.-V. Data curation, Formal Analysis, Validation, Writing – review & editing; L.A.A.-P. Data curation, Formal Analysis, Methodology, Validation, Writing – review & editing. L.B. Data curation, Formal Analysis, Methodology, Validation, Writing – review & editing; G.P.-D. Methodology, Writing – review & editing; O.R.-M. Investigation, Methodology, Resources, Writing – review & editing; J.A.S.-M. Methodology, Resources, Writing – review & editing; G.P.-S. Project administration, Writing – review & editing; D.S.-M. Funding acquisition, Resources, Writing review & editing; W.S. Formal Analysis, Resources, Writing – review & editing; H.E.C.-C. Conceptualization, Formal Analysis, Funding acquisition, Project administration, Supervision, Writing – review & editing; F.C.-P. Investigation, Writing – review & editing; P.P. Formal Analysis, Resources, Writing – review & editing; F.K. Formal Analysis, Resources, Writing – review & editing; A.G.-S. Formal Analysis, Resources, Writing – review & editing; B.L.-D. Conceptualization, Funding acquisition, Resources, Supervision, Validation, Writing – review & editing; C.L.-M. Conceptualization, Data curation, Formal Analysis, Funding acquisition, Investigation, Project administration, Resources, Supervision, Validation, Visualization, Writing – original draft, Writing – review & editing.

## Ethical Responsibilities

The protocol was approved by the ethics, research, and biosafety committees of the Mexican Institute of Social Security (IMSS) National Scientific Research and Ethics Commissions under project numbers R-2020-785-095 and R-2021-785-048. Informed consent was obtained from both patients and vaccinated individuals. This project was conducted in accordance with good clinical practice.

## Protection of Humans and Animals

No animals were used in this protocol, and the protocol was designed to comply with the ethical principles of the Helsinki Declaration, Good Clinical Practices, and the applicable Mexican law.

## Funding

The funding was provided by Laboratorio Avi-Mex, S. A. de C. V. (Avimex) project number R-2021-785-048.

## Acknowledgments

We would like to extend our gratitude to the following individuals for their operational support from Laboratorio Avi-Mex, S. A. de C. V.: Bernardo Lozano Alcántara, Alejandro Ruiz, Rosalba Rodriguez, Leticia Espinosa Gervasio, Rodrigo Yebra Reyes, Vanessa Escamilla Jiménez, Jaime Becerra Jimenez, Lorena Juárez Pedraza, Sandra Yuridia Ang Tinajero, Avelia Ariadna Cuevas Cifuentes, Juan Pablo Robles Alvarez, Avirán Almazán Gutiérrez, Aurora Betsabé Gutiérrez Balderas, Merlenne Rubio Diaz, and Guadalupe Aguilar Rafael. Additionally, we would like to express our appreciation to Dr. Luis Alejandro Sánchez Hurtado from UMAE Hospital de Especialidades, Centro Médico Nacional Siglo XXI, IMSS, Dr. Mauricio Tapia, Dr. Victor Hugo Teposte Miramontes from Centro de Atención Temporal COVID-19 Tlatelolco y Morelos, and Dra. Daira González Rodríguez from Coordinación de Investigación en Salud, IMSS, for their support in obtaining COVID-19 patient samples.

## Data Availability Statement

Individual de-identified participant data will not be shared beyond the limits permitted by the informed consent and Mexican law. Specifically, this includes the sharing of the study protocol, statistical analysis plan, informed consent form, and approved clinical study report. Additionally, other de-identified data allowed under the informed consent and Mexican law may be shared. The data will be made available immediately upon publication and for a duration of 12 months thereafter. Access to the data will be granted solely to investigators with methodologically sound proposals, subject to authorization by an independent review committee and the ethics committees involved in approving the protocol.

### Privacy Rights and Informed Consent

Informed Consent formats were provided and signed by all participants under Good Clinical Practices and Mexican law. Personal data is confidential, and subjects are not identified personally. Their data was analysed and duly de-identified.

## Supplementary Material

**Figure 1S.**
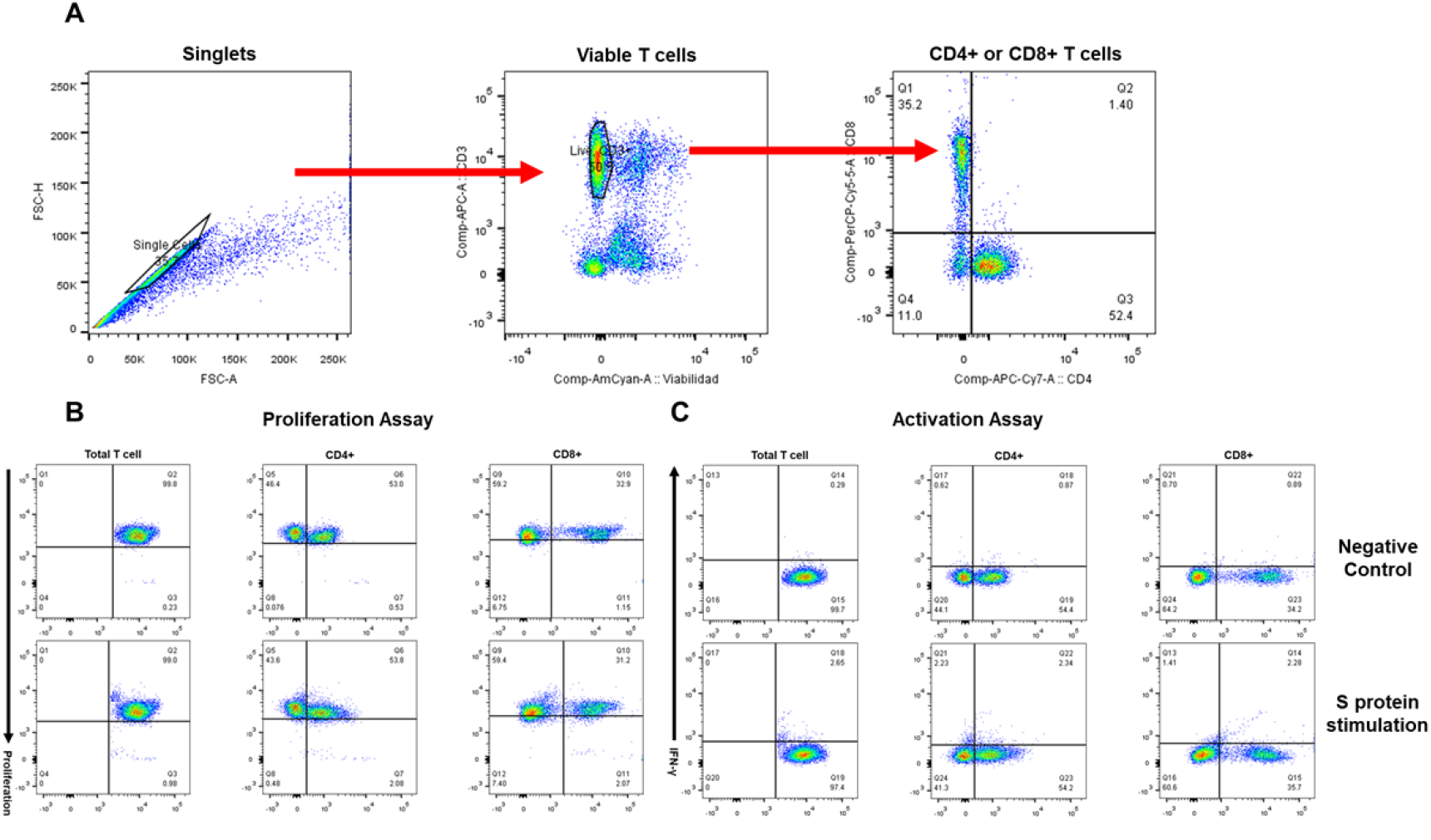
Flow cytometry gating strategy. Representative PBMC sample gating hierarchy. From left to right and top to bottom: (A) FSC-A versus SSC-A for cellular viability and identification of CD4+ or CD8+ T cells, (B) proliferation assay, and (C) individual T cell responses for the production of IFN-γ.

## References

1. WHO Coronavirus (COVID-19) Dashboard | WHO Coronavirus (COVID-19) Dashboard With Vaccination Data. https://covid19.who.int/ [Accessed January 5, 2023]

2. Excess mortality during the Coronavirus pandemic (COVID-19) - Our World in Data. https://ourworldindata.org/excess-mortality-covid [Accessed February 28, 2024]

3. Global Dashboard for Vaccine Equity - UNDP Data Futures Platform. https://data.undp.org/vaccine-equity/ [Accessed January 5, 2023]

4. Hsieh CL, Goldsmith JA, Schaub JM, DiVenere AM, Kuo HC, Javanmardi K, Le KC, Wrapp D, Lee AG, Liu Y, et al. Structure-based design of prefusion-stabilized SARS-CoV-2 spikes. Science (2020) 369:1501–1505. doi: 10.1126/SCIENCE.ABD0826

5. Lara-Puente JH, Carreño JM, Sun W, Suárez-Martínez A, Ramírez-Martínez L, Quezada-Monroy F, de la Rosa GP, Vigueras-Moreno R, Singh G, Rojas-Martínez O, et al. Safety and Immunogenicity of a Newcastle Disease Virus Vector-Based SARS-CoV-2 Vaccine Candidate, AVX/COVID-12-HEXAPRO (Patria), in Pigs. MBio (2021) 12: doi: 10.1128/MBIO.01908-21

6. Sun W, Liu Y, Amanat F, González-Domínguez I, McCroskery S, Slamanig S, Coughlan L, Rosado V, Lemus N, Jangra S, et al. A Newcastle disease virus expressing a stabilized spike protein of SARS-CoV-2 induces protective immune responses. Nat Commun (2021) 12:1–14. doi: 10.1038/s41467-021-26499-y

7. Park JG, Oladunni FS, Rohaim MA, Whittingham-Dowd J, Tollitt J, Hodges MDJ, Fathallah N, Assas MB, Alhazmi W, Almilaibary A, et al. Immunogenicity and protective efficacy of an intranasal live-attenuated vaccine against SARS-CoV-2. iScience (2021) 24:102941. doi: 10.1016/j.isci.2021.102941

8. Ponce-de-León S, Torres M, Soto-Ramírez LE, Calva JJ, Santillán-Doherty P, Carranza-Salazar DE, Carreño JM, Carranza C, Juárez E, Carreto-Binaghi LE, et al. Interim safety and immunogenicity results from an NDV-based COVID-19 vaccine phase I trial in Mexico. npj Vaccines (2023) 8:67. doi: 10.1038/s41541-023-00662-6

9. Choi A, Koch M, Wu K, Chu L, Ma LZ, Hill A, Nunna N, Huang W, Oestreicher J, Colpitts T, et al. Safety and immunogenicity of SARS-CoV-2 variant mRNA vaccine boosters in healthy adults: an interim analysis. Nat Med 2021 2711 (2021) 27:2025–2031. doi: 10.1038/s41591-021-01527-y

10. Guerrera G, Picozza M, D’Orso S, Placido R, Pirronello M, Verdiani A, Termine A, Fabrizio C, Giannessi F, Sambucci M, et al. BNT162b2 vaccination induces durable SARS-CoV-2– specific T cells with a stem cell memory phenotype. Sci Immunol (2023) 6:eabl5344. doi: 10.1126/sciimmunol.abl5344

11. Shirvani E, Samal SK. Newcastle Disease Virus as a Vaccine Vector for SARS-CoV-2. Pathog (Basel, Switzerland) (2020) 9:1–8. doi: 10.3390/PATHOGENS9080619

12. Nakaya T, Cros J, Park M-S, Nakaya Y, Zheng H, Sagrera A, Villar E, Garc I - Sastre A, Palese P. Recombinant Newcastle Disease Virus as a Vaccine Vector. J Virol (2001) 75:11868–11873. doi: 10.1128/jvi.75.23.11868-11873.2001

13. Vigil A, Martinez O, Chua MA, García-Sastre A. Recombinant Newcastle disease virus as a vaccine vector for cancer therapy. Mol Ther (2008) 16:1883–1890. doi: 10.1038/mt.2008.181

14. Cheng H, Peng Z, Luo W, Si S, Mo M, Zhou H, Xin X, Liu H, Yu Y. Efficacy and safety of covid-19 vaccines in phase iii trials: A meta-analysis. Vaccines (2021) 9: doi: 10.3390/VACCINES9060582/S1

15. Hornsby H, Nicols AR, Longet S, Liu C, Tomic A, Angyal A, Kronsteiner B, Tyerman JK, Tipton T, Zhang P, et al. Omicron infection following vaccination enhances a broad spectrum of immune responses dependent on infection history. Nat Commun 2023 141 (2023) 14:1–16. doi: 10.1038/s41467-023-40592-4

16. Altarawneh HN, Chemaitelly H, Ayoub HH, Tang P, Hasan MR, Yassine HM, Al-Khatib HA, Smatti MK, Coyle P, Al-Kanaani Z, et al. Effects of Previous Infection and Vaccination on Symptomatic Omicron Infections. N Engl J Med (2022) 387:21–34. doi: 10.1056/NEJMOA2203965

17. Vogel AB, Kanevsky I, Che Y, Swanson KA, Muik A, Vormehr M, Kranz LM, Walzer KC, Hein S, Güler A, et al. BNT162b vaccines protect rhesus macaques from SARS-CoV-2. Nature (2021) 592:283–289. doi: 10.1038/s41586-021-03275-y

18. Watanabe Y, Mendonça L, Allen ER, Howe A, Lee M, Allen JD, Chawla H, Pulido D, Donnellan F, Davies H, et al. Native-like SARS-CoV-2 Spike Glycoprotein Expressed by ChAdOx1 nCoV-19/AZD1222 Vaccine. ACS Cent Sci (2021) 7:594–602. doi: 10.1021/acscentsci.1c00080

19. Long QX, Liu BZ, Deng HJ, Wu GC, Deng K, Chen YK, Liao P, Qiu JF, Lin Y, Cai XF, et al. Antibody responses to SARS-CoV-2 in patients with COVID-19. Nat Med 2020 266 (2020) 26:845–848. doi: 10.1038/s41591-020-0897-1

20. Bahnan W, Wrighton S, Sundwall M, Bläckberg A, Larsson O, Höglund U, Khakzad H, Godzwon M, Walle M, Elder E, et al. Spike-Dependent Opsonization Indicates Both Dose-Dependent Inhibition of Phagocytosis and That Non-Neutralizing Antibodies Can Confer Protection to SARS-CoV-2. Front Immunol (2022) 12:1–17. doi: 10.3389/fimmu.2021.808932

21. Kaneko N, Kuo HH, Boucau J, Farmer JR, Allard-Chamard H, Mahajan VS, Piechocka-Trocha A, Lefteri K, Osborn M, Bals J, et al. Loss of Bcl-6-Expressing T Follicular Helper Cells and Germinal Centers in COVID-19. Cell (2020) 183:143-157.e13. doi: 10.1016/j.cell.2020.08.025

22. Dugan HL, Stamper CT, Li L, Changrob S, Asby NW, Halfmann PJ, Zheng N-Y, Huang M, Shaw DG, Cobb MS, et al. Profiling B cell immunodominance after SARS-CoV-2 infection reveals antibody evolution to non-neutralizing viral targets. Immunity (2021) doi: 10.1016/j.immuni.2021.05.001

23. Killikelly A, Tunis M, House A, Quach C, Vaudry W, Moore D. Overview of the respiratory syncytial virus vaccine candidate pipeline in Canada. Canada Commun Dis Rep (2020) 46:56–61. doi: 10.14745/ccdr.v46i04a01

24. Hiremath J, Kang K Il, Xia M, Elaish M, Binjawadagi B, Ouyang K, Dhakal S, Arcos J, Torrelles JB, Jiang X, et al. Entrapment of H1N1 influenza virus derived conserved peptides in PLGA nanoparticles enhances T cell response and vaccine efficacy in pigs. PLoS One (2016) 11:1–15. doi: 10.1371/journal.pone.0151922

25. Pacheco-Olvera DL, Saint Remy-Hernández S, García-Valeriano MG, Rivera-Hernández T, López-Macías C. Bioinformatic Analysis of B- and T-cell Epitopes from SARS-CoV-2 Structural Proteins and their Potential Cross-reactivity with Emerging Variants and other Human Coronaviruses. Arch Med Res (2022) 53:694–710. doi: 10.1016/J.ARCMED.2022.10.007

26. Meyer S, Blaas I, Bollineni RC, Delic-Sarac M, Tran TT, Knetter C, Dai KZ, Madssen TS, Vaage JT, Gustavsen A, et al. Prevalent and immunodominant CD8 T cell epitopes are conserved in SARS-CoV-2 variants. Cell Rep (2023) 42: doi: 10.1016/J.CELREP.2023.111995

27. Singh G, Abbad A, Kleiner G, Srivastava K, Gleason C, Carreño JM, Simon V, Krammer F, Andre D, Bermúdez-González MC, et al. The post-COVID-19 population has a high prevalence of cross-reactive antibodies to spikes from all Orthocoronavirinae genera. MBio (2024) doi: 10.1128/MBIO.02250-23

